## Supplemental data for "Sensory kinase KdpD is a tandem serine histidine kinase controlling K^+^ pump KdpFABC on the translational and post-transcriptional level"

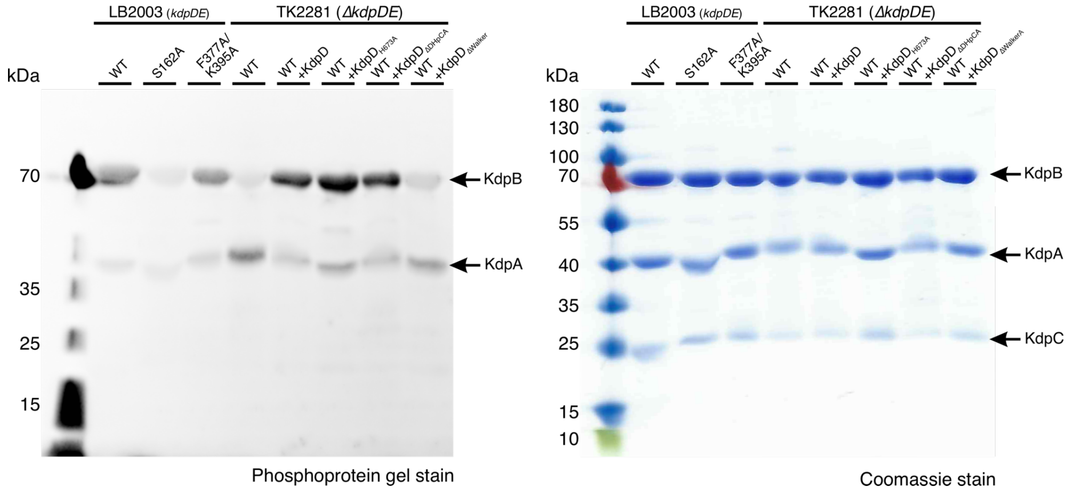


**Supplementary Figure 1: KdpD phosphorylates $KdpB_{S162}$ via a Walker A motif in the KdpD domain – full phosphoprotein and Coomassie stains.** After cultivation at high external $K^+$ concentrations, KdpFABC constructs were purified from *E. coli* LB2003 cells, which natively express *kdpDE*, or from *E. coli* TK2281 cells, in which the complete *kdpFABCDE* operon is deleted. In the latter setting, KdpD variants were reintroduced as indicated by expression from an orthogonal plasmid. **A**, Phosphoprotein gel stain of purified KdpFABC separated by SDS-PAGE, indicating the phosphorylation state of $KdpB_{S162}$. An additional signal corresponding to KdpA likely is caused by a tightly associated cardiolipin molecule (Silberberg et al., 2021). **B**, The Coomassie stain indicates the comparable amount of protein loaded per lane. Protein bands corresponding to KdpA, KdpB, and KdpC are indicated.

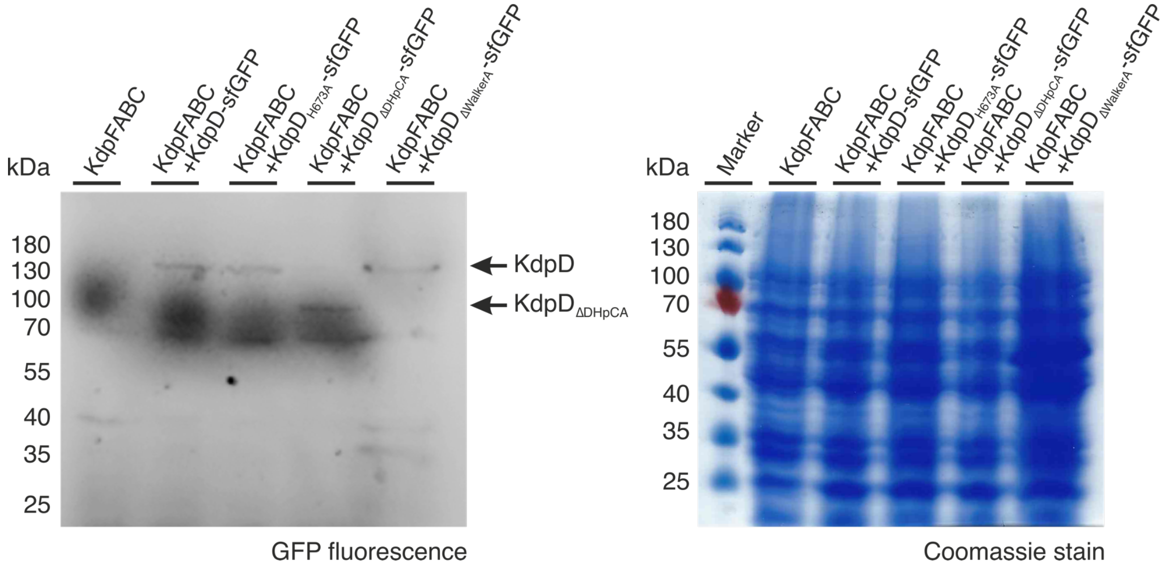


**Supplementary Figure 2: Co-expression of WT KdpFABC with KdpD constructs in *E. coli* TK2281 cells.** Whole-cell samples of *E. coli* TK2281 cells co-transformed with plasmids encoding for WT KdpFABC and different KdpD variants, the latter C-terminally fused to sfGFP, were separated by SDS-PAGE to show that KdpD constructs were successfully produced. **A**, GFP fluorescence (ex. 485 nm, em. 535 nm) at molecular weights corresponding to KdpD-sfGFP (124 kDa) or $KdpD_{\Delta DHpCA}$-sfGFP (100 kDa) indicates comparable protein levels of the different KdpD constructs. **B**, The Coomassie stain indicates the comparable amount of cell lysate loaded per lane.

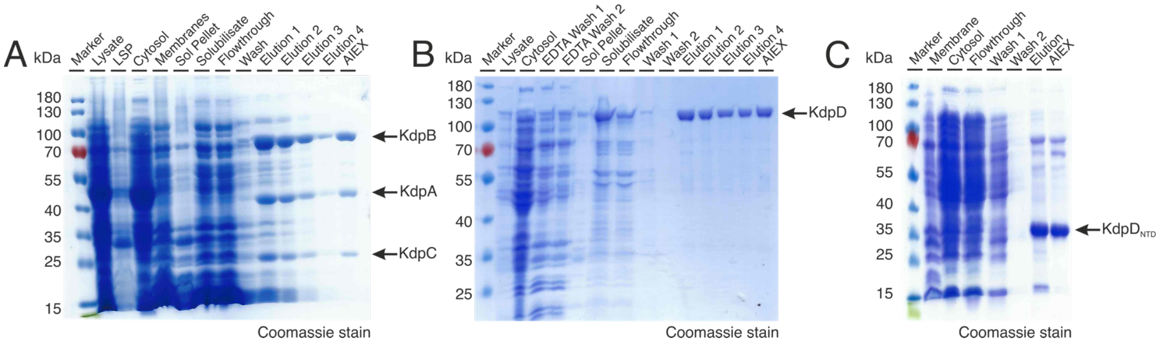


**Supplementary Figure 3: Purification of KdpFAB$_{D307N}$C, KdpD, and KdpD$_{NTD}$ from *E. coli* TK2281 cells.** SDS-PAGE showing steps from the purification of KdpFAB$_{D307N}$C-His$_{10}$ (**A**), KdpD-His$_{10}$ (**B**), or KdpD$_{NTD}$-His$_{10}$ (**C**) from *E. coli* TK2281 cells, with a high degree of purity after Ni$^{2+}$-NTA and AIEX chromatography. Abbreviations: Low-speed pellet (LSP), Solubilization pellet (Sol Pellet); Flowthrough, Wash, and Elutions from Ni$^{2+}$-NTA column. AIEX eluted with increasing NaCl concentration from a HiTrap Q HP column (Cytiva; Marlborough, MA, USA).

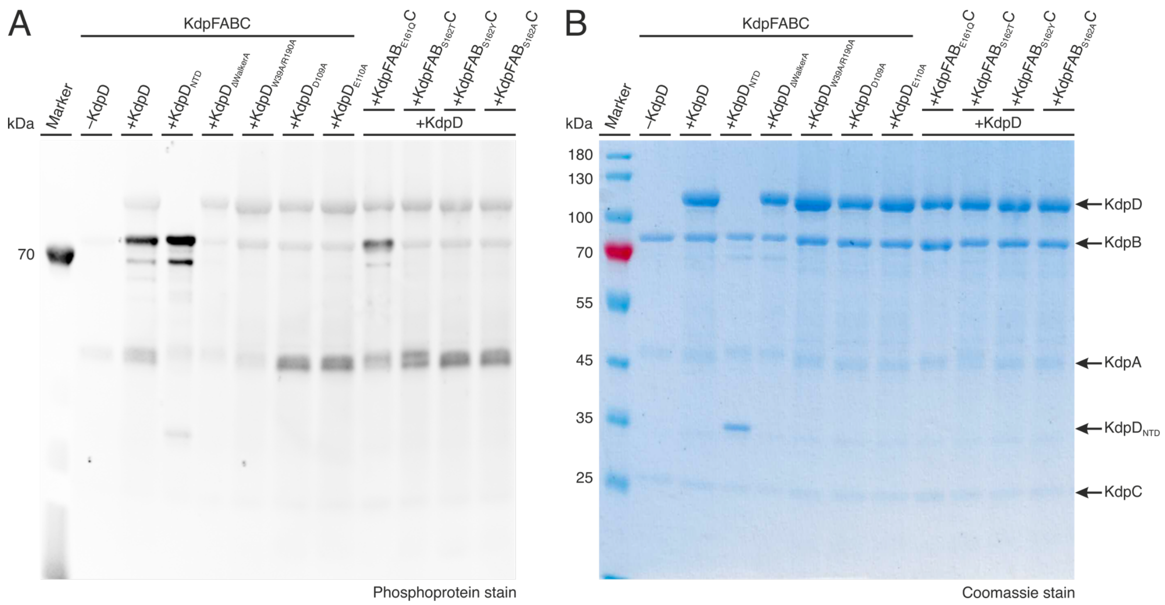


**Supplementary Figure 4: Phosphorylation of KdpB$_{S162}$ by KdpD – full phosphoprotein and Coomassie stains.** KdpFAB$_{D307N}$C and KdpD constructs produced in *E. coli* TK2281 cells were purified separately and mixed in vitro in the presence of 400 mM KCl and 5 mM ATP. KdpB phosphorylation levels after 30 min were analyzed by SDS-PAGE and subsequent phosphoprotein gel stain. **A**, Phosphoprotein gel stain of KdpFABC with different KdpD constructs, separated by SDS-PAGE, indicating the phosphorylation state of KdpB$_{S162}$. **B**, The Coomassie stain indicates the comparable amount of protein loaded per lane. Protein bands corresponding to KdpA, KdpB, KdpC, KdpD and KdpD$_{NTD}$ are indicated.

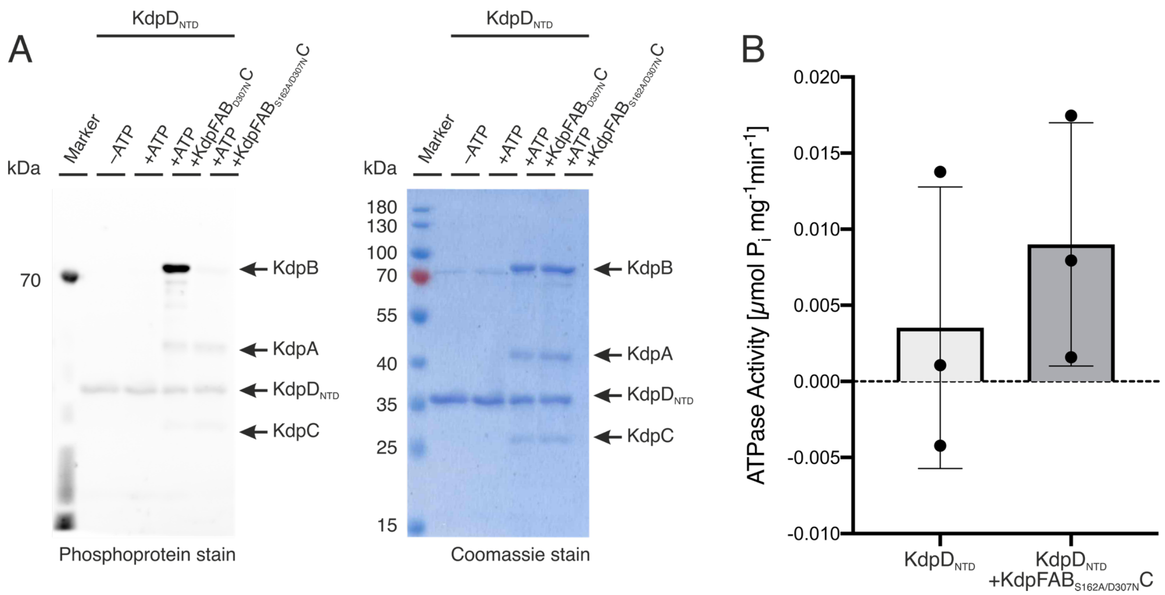


**Supplementary Figure 5: KdpD$_{NTD}$ functions without autophosphorylation or ATP hydrolysis. A**, The phosphorylation state of KdpD$_{NTD}$ was analyzed by phosphoprotein stain in the absence and presence of 5 mM ATP, the substrate KdpFAB$_{D307N}$C, and the substrate with a mutated phosphate acceptor KdpFAB$_{S162A/D307N}$C. The corresponding Coomassie stain indicates sufficient purity and similar amounts of protein loaded. **B**, ATPase activity of purified KdpD$_{NTD}$ in the absence and presence of KdpFAB$_{S162A/D307N}$C, indicating no ATP hydrolysis by the kinase.

A

| | | | | | | | |
|---|---|---|---|---|---|---|---|
| Acidobacteriota | Actinomycetota | Aquificota | Archaea | Armatimonadota | Bacillota | Bacteroidota | Balneolota |
| 37 | 1511 | 3 | 49 | 5 | 694 | 583 | 1 |
| Bdellovibrionota | Campylobacterota | Chlamydiota | Chlorobiota | Chloroflexota | Cyanobacteriota | Deferribacterota | Deinococcota |
| 10 | 19 | 6 | 3 | 25 | 131 | 1 | 28 |
| Desulfobacterota | Fusobacteriota | Gemmatimonadota | Lentisphaerota | Mycoplasmatota | Myxococcota | Nitrospirota | Planctomycetota |
| 10 | 5 | 3 | 2 | 2 | 35 | 10 | 31 |
| Pseudomonadota | Spirochaetota | Synergistota | Thermodesulfobacteriota | Thermodesulfobiota | Thermomicrobiota | Thermotogota | Verrucomicrobiota |
| 2161 | 57 | 1 | 35 | 2 | 1 | 2 | 32 |

B

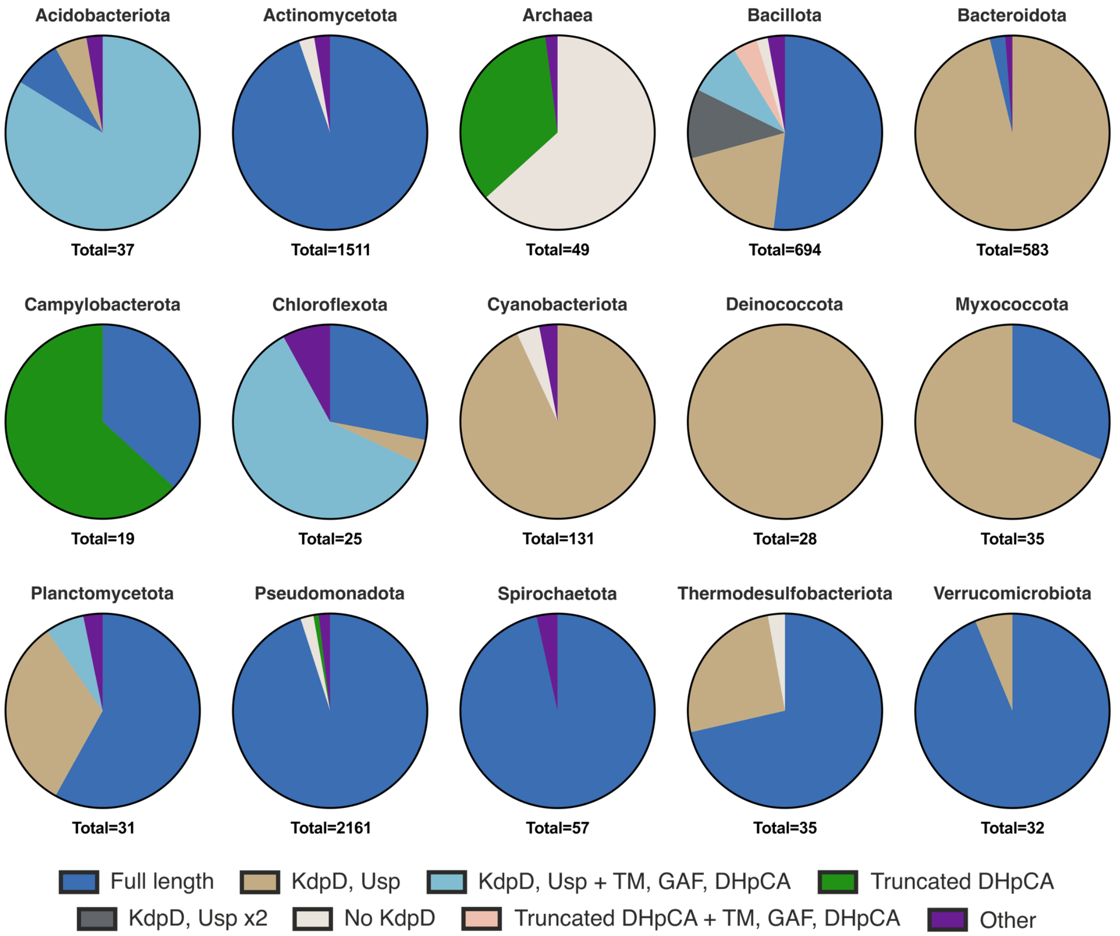


**Supplementary Figure 6: KdpD versions in different phyla.** KdpD sequences from 5495 species featuring an annotation for the Kdp pump in the UniProt database were extracted. The dataset was based on that of a previous bioinformatic study (Wang et al., 2019), which was expanded to include all other species from the LPSN database fitting the parameters (Parte et al., 2020). **A**, Number of species analyzed per phylum. **B**, Distribution of KdpD versions in the 15 largest phylogenic groups of the dataset.

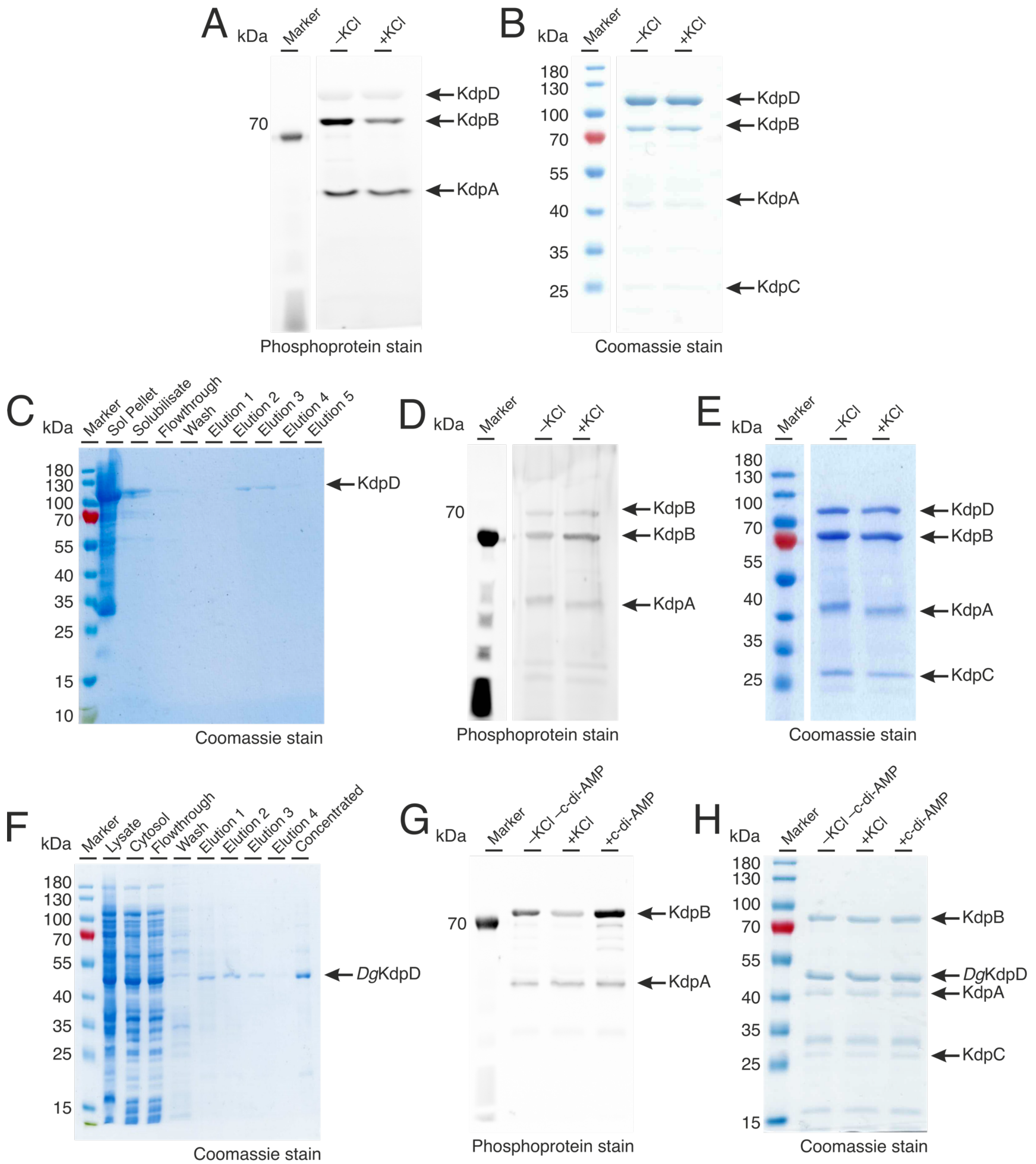


**Supplementary Figure 7: Stimulation of KdpD ASK activity – full phosphoprotein and Coomassie stains and purifications of *E. coli* KdpD in SMALPs and *D. geothermalis* KdpD.** KdpFAB$_{D307N}$C and KdpD constructs produced in *E. coli* TK2281 cells were purified separately and mixed in vitro with 5 mM ATP in the absence and presence of 400 mM KCl or 1 mM c-di-AMP. KdpB phosphorylation levels after 30 min were analyzed by SDS-PAGE and subsequent phosphoprotein gel stain. **A**, **B**, Phosphoprotein- and Coomassie-stained SDS-PAGE showing the ASK activity of DDM-solubilized *E. coli* KdpD in the absence and presence of 400 mM KCl. **C**, Purification of *E. coli* KdpD solubilized in SMALPs from *E. coli* TK2281 cells, with a high degree of purity after Ni$^{2+}$-NTA chromatography. **D**, **E,** Phosphoprotein- and Coomassie-stained SDS-PAGE showing stimulation of ASK activity of SMALP-solubilized *E. coli* KdpD by 400 mM KCl. **E**, Purification of *D. geothermalis* KdpD from *E. coli* TK2281 cells, with a high degree of purity after Ni$^{2+}$-NTA chromatography. **F**, **G**, Phosphoprotein- and Coomassie-stained SDS-PAGE showing the ASK activity of *D. geothermalis* KdpD in the absence and presence of 400 mM KCl or 1 mM c-di-AMP.
