## Supplementary material for "Sensory kinase KdpD is a tandem serine histidine kinase controlling K^+^ pump KdpFABC on the translational and post-transcriptional level": Table S1

**Table S1: Cloning Primers**

| Name | Source | Sequence |
|---|---|---|
| kdpD_FX_for | Eurofins Genomics | ATATATGCTCTTCTAGTAATAACGAACCCTTACGTCCCGACCCCGATCG |
| kdpD_FX_rev | Eurofins Genomics | TATATAGCTCTTCCTGCCATATCCTCATGAAATTCTTCAAGTTC |
| kdpD_NTD_FX_rev | Eurofins Genomics | TATATAGCTCTTCATGCCCACTTGTCTTTAAATGAGCGGTTATCCGGCG |
| kdpD_H673A_for | Eurofins Genomics | GGCGGCGCTTTCGGCAGATTTACGCACGC |
| kdpD_H673A_rev | Eurofins Genomics | GCGTGCGTAAATCTGCCGAAAGCGCCGCC |
| kdpD_G37A_K38A_T39C_for | Eurofins Genomics | GTGCAGGCGTCGCAGCATGCTGGGCGATGCTGGCAGAAGC |
| kdpD_G37A_K38A_T39C_rev | Eurofins Genomics | GCATCGCCCAGCATGCTGCGACGCCTGCACAGGCACCG |
| kdpD_W39A_for | Eurofins Genomics | CGTCGGGAAGACCGCTGCGATGCTGGCAG |
| kdpD_W39A_rev | Eurofins Genomics | CTGCCAGCATCGCAGCGGTCTTCCCGACG |
| kdpD_R190A_for | Eurofins Genomics | CGATCTGCGCCAGGCTCTGAAAGAAGGC |
| kdpD_R190A_rev | Eurofins Genomics | GCCTTCTTTCAGAGCCTGGCGCAGATCG |
| kdpD_D109A_for | Eurofins Genomics | GGCGCTGATCTTAATGGCTGAACTGGCGCACAGTAATGC |
| kdpD_D109A_rev | Eurofins Genomics | GCATTACTGTGCGCCAGTTCAGCCATTAAGATCAGCGCC |
| kdpD_E110A_for | Eurofins Genomics | CGCTGATCTTAATGGACGCACTGGCGCACAGTAATGCG |
| kdpD_E110A_rev | Eurofins Genomics | CGCATTACTGTGCGCCAGTGCGTCCATTAAGATCAGCG |
| kdpD_XbaI_for | Eurofins Genomics | ATATATTCTAGAATGAATAACGAACCCTTACGTCC |
| KdpD_PstI_rev | Eurofins Genomics | TATATACTGCAGTCACATATCCTCATGAAATTCTTCAAG |
| kdpD_dDHpCA_PstI_rev | Eurofins Genomics | ATATATCTGCAGTCACTGTTCACGTTCGCTTGCC |
| PstI-pBAD33_for | Eurofins Genomics | ATATATCTGCAGGCATGCAAGCTTGG |
| kdpD_PstI_noStop_rev | Eurofins Genomics | TATATACTGCAGCATATCCTCATGAAATTCTTCAAGTTCAGGG |
| kdpD_dDHpCA_NoStop_PstI_rev | Eurofins Genomics | TATATACTGCAGCTGTTCACGTTCGCTTGCC |
| PstI_sfGFP_for | Eurofins Genomics | ATATATCTGCAGAGCAAAGGAGAAGAACTTTTCACTGG |
| His10_HindIII_rev | Eurofins Genomics | ATATATAAGCTTTTAATGATGGTGATGATGATGGTGATG |
| KdpB_E161Q_for | Eurofins Genomics | CGCCATCACCGGACAATCGGCACC |
| KdpB_E161Q_rev | Eurofins Genomics | GGTGCCGATTGTCCGGTGATGGCG |
| kdpB_S162A_for | Eurofins Genomics | GCGCCATCACCGGGGAAGCGGCACC |
| kdpB_S162A_rev | Eurofins Genomics | ACCGGTGCCGCTTCCCCGGTGATGG |
| kdpB_S162T_for | Eurofins Genomics | CCATCACCGGGGAAACCGCACCAGTGATCCGTGAATCC |
| kdpB_S162T_rev | Eurofins Genomics | CGGATCACTGGTGCGGTTTCCCCGGTGATGG |
| kdpB_S162Y_for | Eurofins Genomics | CCATCACCGGGGAATATGCACCAGTGATCCGTGAATCC |

| | | |
|---|---|---|
| kdpB_S162Y_rev | Eurofins Genomics | GGATTCACGGATCACTGGTGCATATTCCC CGGTGATGG |
| kdpB_D307N_for | Eurofins Genomics | GCTACTGAATAAAACCGGCACCATCAC |
| kdpB_D307N_rev | Eurofins Genomics | GGTGCCGGTTTTATTCAGTAGCAGAACG |
| kdpB_F377A_for | Eurofins Genomics | CCTTTGTACCGGCAACTGCGCAAAGC |
| kdpB_F377A_rev | Eurofins Genomics | GCTTTGCGCAGTTGCCGGTACAAAGG |
| kdpB_K395A_for | Eurofins Genomics | CATGATCCGTGCAGGTTCTGTCGATGCC |
| kdpB_K395A_rev | Eurofins Genomics | CGACAGAACCTGCACGGATCATGCGG |
| kdpD_dg_FX_for | Eurofins Genomics | ATATATGCTCTTCTAGTCCTGGTCCTACCC GCCTGAATCCGCC |
| kdpD_dg_FX_rev | Eurofins Genomics | TATATAGCTCTTCATGCATCCCGACTGATG ACGTAGACATCCAC |
